# Nucleolar TFIIE plays a role in ribosomal biogenesis and performance

**DOI:** 10.1101/2020.12.16.423046

**Authors:** Tamara Phan, Pallab Maity, Christina Ludwig, Lisa Streit, Jens Michaelis, Karin Scharffetter-Kochanek, Sebastian Iben

## Abstract

Ribosome biogenesis is a highly energy-demanding process in eukaryotes which requires the concerted action of all three RNA polymerases. In RNA polymerase II transcription, the general transcription factor TFIIH is recruited by TFIIE to the initiation site of protein-coding genes. Distinct mutations in TFIIH and TFIIE give rise to the degenerative disorder trichothiodystrophy (TTD). Here we uncovered an unexpected role of TFIIE in ribosomal RNA synthesis by RNA polymerase I. With high resolution microscopy we detected TFIIE in the nucleolus where TFIIE binds to actively transcribed rDNA. Mutations in TFIIE affects gene-occupancy of RNA polymerase I, rRNA maturation, ribosomal assembly and performance. In consequence, the elevated translational error rate with imbalanced protein synthesis and turnover results in an increase in heat-sensitive proteins. Collectively, mutations in TFIIE – due to impaired ribosomal biogenesis and translational accuracy – lead to a loss of protein homeostasis (proteostasis) which can partly explain the clinical phenotype in TTD.

## Introduction

The biogenesis of ribosomes starts with the synthesis of pre-ribosomal RNA (rRNA) by RNA polymerase I. The pre-rRNA is co-transcriptionally matured, processed and assembled with ribosomal proteins (RP) to pre-ribosomal particles within the nucleolus (Bassler & Hurt, 2019). Ribosomal biogenesis requires the concerted action of all three RNA polymerases. While RNA polymerase I and III provide the rRNAs for ribosomal biogenesis (60% of total transcription), 50% of RNA polymerase II transcription activity is devoted to the production of mRNAs coding for ribosomal proteins (Warner, 1999). All RNA polymerases are dependent on unique sets of basal transcription factors that recognize the core promoter, recruit the RNA polymerases and organize the first steps of productive transcription (Roeder & Rutter, 1969, Vannini & Cramer, 2012, Zhang, Najmi et al., 2017). Some of these basal transcription factors are shared between the RNA polymerases like TBP, that, decorated with class-specific associated proteins, is essential for all three RNA polymerases (Cormack & Struhl, 1992, Schultz, Reeder et al., 1992). The DNA-repair factor TFIIH, consisting of ten subunits, is a basal transcription factor of RNA polymerase II. TFIIH melts the promoter and contributes to promoter escape of the polymerase (Rimel & Taatjes, 2018). TFIIH is also essential for RNA polymerase I transcription (Iben, Tschochner et al., 2002) as an elongation factor (Assfalg, Lebedev et al., 2012, Nonnekens, Perez-Fernandez et al., 2013). Mutations in TFIIH cause the photosensitive form of the multisystem disorder trichothiodystrophy (TTD), a syndrome with delayed development, microcephaly, neurological degeneration, recurrent, infections and eponymous skin and hair abnormalities. TTD is mainly attributed to a transcription syndrome (Compe, Genes et al., 2019, Theil, Mandemaker et al., 2017). However, recently discovered mutations in aminoacyl-tRNA synthetases that result in TTD may shift this classification towards a “gene expression syndrome” referred to a disturbed protein translation at the ribosome (Kuo, Theil et al., 2019, Theil, Botta et al., 2019).

TFIIH mutations can also cause the premature aging disease Cockayne syndrome (CS) that resembles TTD with symptoms of developmental delay, microcephaly and neurodegeneration. Earlier TFIIH mutations have been reported to affect RNA polymerase I transcription and the processing of the pre-rRNA in cellular and mouse models of CS and TTD (Assfalg et al., 2012, Nonnekens et al., 2013). Interestingly, a recent study addressed cellular consequences of CS-mutations for RNA polymerase I transcription and uncovered disturbed RNA polymerase I transcription to be responsible for the generation of highly defective ribosomes. In the same report, defective ribosomes were found to display a profoundly reduced translational accuracy giving rise to a loss of proteostasis that in turn represses RNA polymerase I transcription (Alupei, Maity et al., 2018). As mutations in subunits of RNA polymerase I or the basal transcription factor UBF were identified in syndromes of childhood neurodegeneration (Edvardson, Nicolae et al., 2017, Kara, Koroglu et al., 2017, Sedlackova, Lassuthova et al., 2019), it is tempting to speculate that a disturbed RNA polymerase I transcription might be causally involved in the neurodegeneration of CS and TTD.

A subset of TTD cases is caused by mutations in the RNA polymerase II transcription factor TFIIE, respective the β-subunit of this heterodimeric factor (Grunberg, Warfield et al., 2012, Kuschal, Botta et al., 2016, Theil et al., 2017). During the last steps of transcription initiation of RNA polymerase II, TFIIEβ binds to the preinitiation complex (PIC) and recruits TFIIEα and TFIIH to the initiation site. After phosphorylation of RNAP2 by TFIIH, TFIIEα is released from the PIC before DNA opening, while TFIIEβ dissociates after DNA opening (Compe et al., 2019, Grunberg et al., 2012).

We here hypothesize, that TTD is a syndrome with a pathophysiologic involvement of ribosomal biogenesis and protein homeostasis. If this is the case, TFIIEβ should be involved in these central processes as its mutation provokes the same TTD disease.

Here we discovered an unanticipated role of TFIIE in rRNA synthesis, processing and performance. We detected TFIIE in the nucleolus and showed TFIIE binding to active transcribed rDNA. TFIIEβ-deficient TTD cells display reduced RNA polymerase I binding to the rDNA and disturbed pre-rRNA maturation. Impaired rRNA maturation subsequently leads to reduced ribosomal stability and unbalanced ribosomal composition in NP-TTD cells. As a consequence, ribosomes of TFIIEβ-deficient TTD cells show increased translational inaccuracy and an imbalanced protein homeostasis

## Results

### TFIIE is enriched in transcriptionally active nucleoli and binds to ribosomal DNA

TFIIE mutations cause TTD, a disease mainly provoked by mutations in the multifunctional TFIIH complex. As the TFIIH complex plays an essential role in RNA polymerase I transcription (Iben et al., 2002), we addressed the question if TFIIE is also involved in ribosomal biogenesis. RNA polymerase I transcription takes place in the nucleolus, the densest cellular organelle. To investigate if TFIIE localizes to the nucleolus, co-localization studies employing confocal microscopy with RNA polymerase I were performed and further refined by high resolution STED microscopy (Figure 1A, Supplement Figure S1A). We observed a distinct enrichment of TFIIEβ in the nucleoli and a close co-localization with RNA polymerase I. The large subunit of TFIIE, TFIIEα showed a similar pattern when co-stained with RNA polymerase I. This data indicate that both subunits of TFIIE are enriched in the nucleoli. Remarkably, this enrichment is lost when RNA polymerase I transcription is blocked, by the specific inhibitor CX5461 leading to a re-organization of RNA polymerase I in nucleolar caps (Figure 1B and C, Supplement Figure S1B). Interestingly, TFIIF, a general transcription factor of RNA polymerase II and one interaction partner of TFIIE, is not enriched in the nucleolus (Supplement Figure S1C), indicating a RNA polymerase II-independent localization of TFIIE in the nucleolus. As the TFIIE complex binds to the promoter DNA of RNA polymerase II genes (Compe et al., 2019, Grunberg et al., 2012) we further explored a potential binding of TFIIE to the ribosomal DNA using ChIP experiments. In fact, TFIIE binding to the rDNA was to the end of the coding region of the 28SrRNA was detected, while TFIIE did not bind to the promoter of the rDNA. A specific binding of both subunits of TFIIE to the rDNA is clearly detectable by semi-quantitative and quantitative PCR (Figures 1D, E left side, Supplement Figure S1D-E). These findings are supported by further experiments depicted in Figure 1D and E (right panels). In fact, using ChIP-re-ChIP experiments we show that both TFIIE subunits were precipitated with RNA polymerase I antibodies. These data imply that TFIIE and RNA polymerase I bind to the same rDNA molecules constituting the transcriptionally active fraction of the rDNA. This rDNA binding is reduced for TFIIEβ and RNA polymerase I, when treating the cells with CX5461 (Figure 1F). Thus, TFIIE localizes to the nucleolus and associates with transcriptional active rDNA molecules providing evidence for a possible role of TFIIE in ribosomal biogenesis.

**Figure 1.**
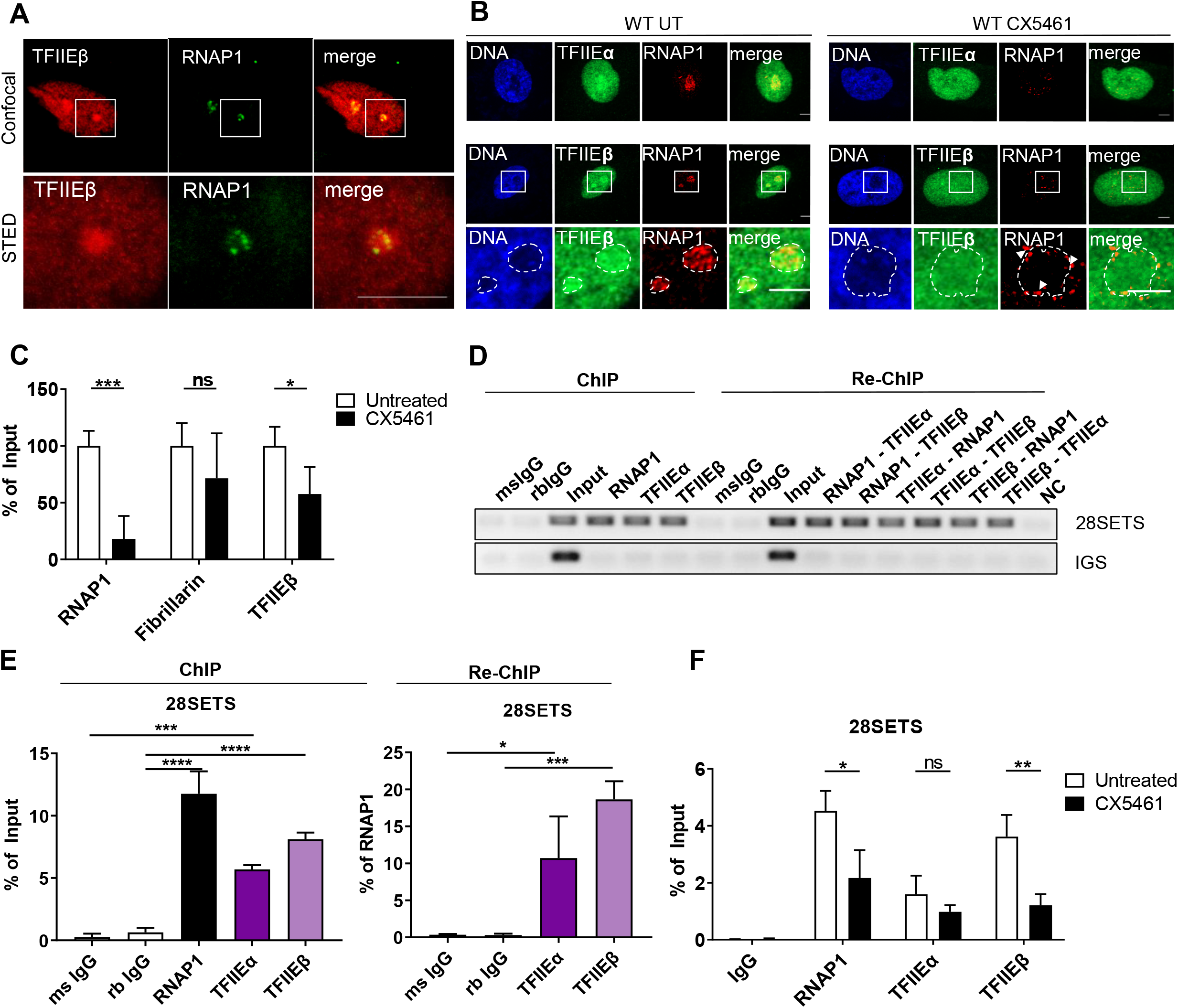
Both TFIIE subunits localize to the nucleolus and associate with rDNA genes. **(A)** High resolution STED microscopy images showing co-localization of TFIIEβ with RNAP1. More STED images of TFIIEβ with Fibrillarin or Nucleolin are given in the supplement Figure S1A. Scale bar 5 µm. **(B)** Confocal immunofluorescence microscopy (100X) of WT cells indicates co-localization of both TFIIE subunits with RNAP1. Inhibition of RNA polymerase I transcription in WT cells by 1 µM CX5461 result in a re-distribution of both TFIIE from the nucleolar enrichment and an organization of RNAP1 to nucleolar caps (arrows) More confocal microscopy images showing co-localization of TFIIEβ with Fibrillarin are given in the supplement Figure S1B. Scale bar 5 µm. **(C)** Quantification of CTCF (corrected total cell fluorescence) of IF performed in (B) indicating reduced TFIIEβ staining in the nucleolus after CX5461 treatment. **(D)** Semi-quantitative PCR analysis of ChIP and sequential ChIP (Re-ChIP) of TFIIE subunits and RNAP1 shows binding of TFIIEβ to the end of the 28S rDNA region (28SETS), and on the same rDNA molecules as RNAP1. Semi-quantitative PCR analysis of the ribosomal intergenic spacer (IGS) region serves as a negative control. **(E)** Quantitative qPCR analysis of ChIP performed in (D) and sequential ChIP performed with RNAP1 indicates significant binding of TFIIE subunits to the same rDNA molecule as RNAP1. More semi-quantitative and qualitative ChIP analysis showing binding of TFIIEβ to the middle of the 28S rDNA region are given in the supplement Figure S1D and S1 E. Data are represented as mean ± SD. ns p > 0.05, * p ≤ 0.05, ** p ≤ 0.01, *** p ≤ 0.001.

### Mutations in TFIIEβ affect rDNA transcription and processing

The two fibroblasts cell line TTD218UT and TTD28N used in this study were isolated from two different NP-TTD patients bearing the same homozygote mutation in D187Y (Supplement Figure S2A,(Kuschal et al., 2016, Theil et al., 2017). Both patient cell lines were reconstituted with a wild-type TFIIEβ fused to GFP: 218UT TFIIEβ and 28N TFIIEβ. For the following experiments the patient cell lines were compared to a healthy control (WT) and the respective reconstituted cell line.

As previously described (Theil et al., 2017), employing western blot analysis we confirmed reduced protein concentration of TFIIEβ in both TTD cell lines indicating that the mutation affects TFIIEβ protein stability. By contrast, protein levels of TFIIH subunits are stable (Supplement Figure S2C-E). As reported, the abundance of TFIIEα protein is diminished although there are no mutations. These findings were confirmed by immunofluorescence stainings of both TFIIE subunits in TTD cells compared to reconstituted cell and WT cell lines (Supplement Figure S2B). TFIIEβ mutations have no impact on rDNA promoter methylation as shown by the methylation restriction enzyme sensitivity assay (Supplement Figure S1F).

To investigate if the reduced TFIIE levels in TTD cells may impact on RNA polymerase I transcription, we examined gene occupancy of the rDNA by RNA polymerase I in more detail. ChIP experiments and the analysis of 16 rDNA regions (Figure 2A) by qPCR maps the density of RNA polymerase I on the ribosomal DNA template and creates a binding profile. The binding profile was then compared between wild-type, reconstituted and TFIIEβ mutated cell lines. As depicted in Figure 2B, the rDNA binding profile of RNA polymerase I is significantly altered in both TTD patient cell lines, with a significant reduction of rDNA binding by RNA polymerase I throughout the entire 28S rDNA. These results imply that the abundance of TFIIE on the rDNA is essential for the proper association of the polymerase with the template.

**Figure 2.**
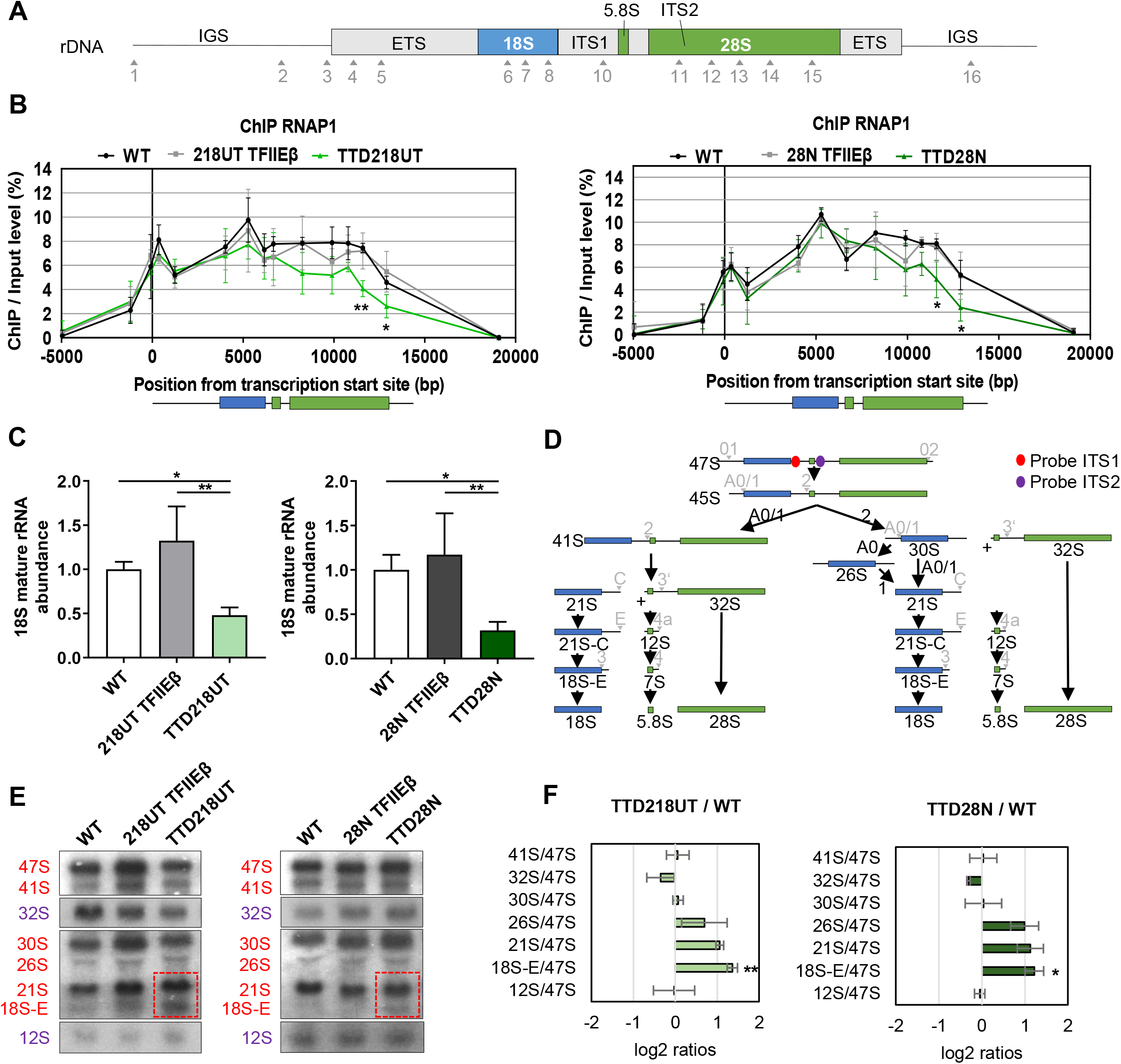
TFIIE influences RNA polymerase I gene occupancy and rRNA processing. **(A)** Schema of the human rDNA unit. The non-coding region ETS, ITS1, and ITS2 (gray) and coding region 18S (blue), 5.8S and 28S (both in green) are illustrated. Primer used for ChIP analysis are indicated by arrowheads and numbers. **(B)** Quantitative qPCR ChIP analysis of RNAP1 in WT, reconstituted and TTD cells indicates reduced binding of RNAP1 to the end of the 28S rRNA coding rDNA (region 14 and 15) in TTD cells. **(C)** Quantitative qPCR analysis indicates reduced abundance of mature 18S rRNA. Quantitative qPCR analysis of pre-courser rRNA are given in the supplement Figure S3 A. **(D)** Schema of the rRNA processing pathway in human cells; adapted from (Mullineux & Lafontaine, 2012). ITS1 and ITS2 probes positions are marked in red and purple, respectively. **(E)** Northern Blot of WT, reconstituted and TTD cells shows accumulation of 21S and 18S-E level in TTD cells. Red and purple color code indicates rRNA species detected by ITS1 and ITS2 probe, respectively. Full images of the probed membrane are given in the supplement Figure S3 B **(F)** Analysis of the northern blots in (E) are displayed as Ratio Analysis of Multiple Precursors (RAMP) profiles;(Wang & Pestov, 2016). Data are represented as mean ± SD. * p ≤ 0.05, ** p ≤ 0.01.

Since TFIIEβ mutations in TTD cells reduce the binding of RNAP1 to the end of the rDNA, we further assessed pre-rRNA levels and mature rRNA abundance by qPCR analysis. No significant reduction of pre-rRNAs was detected (Supplement Figure S3A), however, the amount of mature 18S rRNA was significantly reduced in TTD cells (Figure 2C). The pre-rRNA precursor is co-transcriptionally processed, matured and assembled in pre-ribosomal particles (Bassler & Hurt, 2019). Therefore, and stimulated by previous work (Nonnekens et al., 2013), we were interested to monitor the processing steps from the pre-rRNA to the mature product by Northern blot analysis. In Figure 2D the processing steps are schematically depicted and the position of the probes used to perform Northern blots are indicated. This technique allows to visualize and quantitate all processing intermediates within the rRNA maturation pathway. Northern blot analysis and RAMP profile quantification (Wang & Pestov, 2016) revealed an increased tendency to the accumulation of 21S and a significant accumulation of 18S-E processing intermediate in TTD cells as opposed to WT and reconstituted cells (Figure 2E and F, Supplement Figure S3B-C). Taken together these results show, that mutation in TFIIEβ weakens the interaction of RNA polymerase I with the template that is followed by the accumulation of processing intermediates.

### TFIIEβ mutation affects ribosomal stability and performance

Next, we asked the question if and how disturbed ribosomal biogenesis affects ribosomal composition and the translation process. While mature 18S rRNA is the structural and functional backbone of the small ribosomal subunit 40S, the 5.8S, 28S rRNA, and 5S (a product of RNA polymerase III) are components of the large ribosomal subunit 60S. Since our TTD cells show reduced mature 18S rRNA abundance due to disturbances in rRNA processing, we isolated ribosomes from reconstituted and TTD cells after sucrose gradient centrifugation (Penzo, Carnicelli et al., 2016) and performed quantitative analysis of the ribosomal composition by mass spectrometry (Figure 3A). Although not reaching significance, all detectable ribosomal proteins of the small subunit 40S are reduced in TTD ribosomes compared to the reconstituted cell line. This finding is in line with the reduced presence of mature 18S rRNA in TTD cells. To confirm our mass spectrometry data, we picked three ribosomal proteins of the big and small ribosomal subunit and analyzed their protein level in whole-cell lysate and in purified ribosomes. We were able to visualize a relative underrepresentation of proteins of the small ribosomal subunit in isolated ribosomes (Figure 3B, Supplement Figure S3D-E).

**Figure 3.**
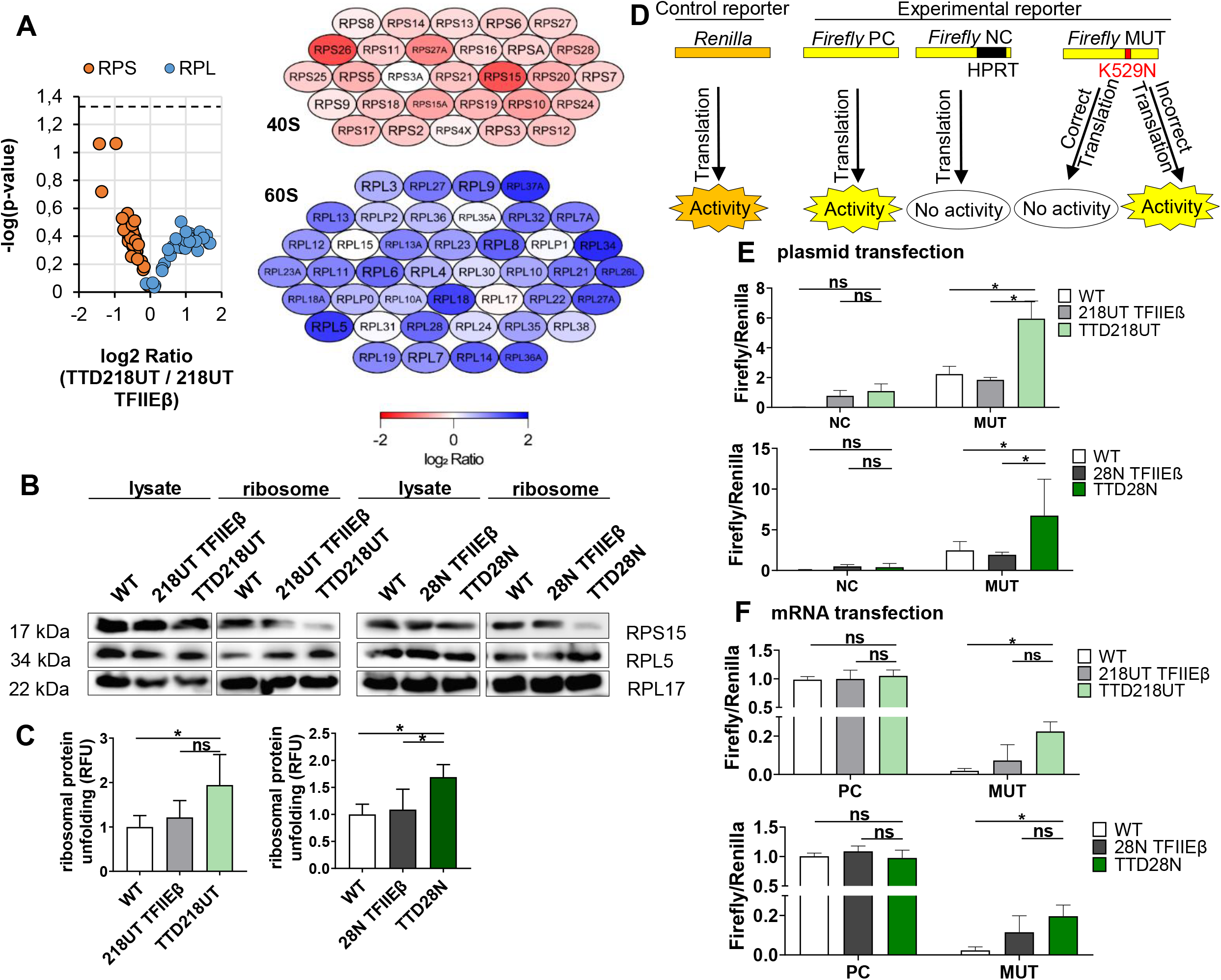
Mutation in TFIIE disturbs ribosomal composition and performance. **(A) Left:** Volcano plot shows a non-significant decreased amount of ribosomal protein of the small 40S ribosome subunit (RPS) and a non-significant increased amount of ribosome protein of the big 60S ribosome subunit (RPL) in TTD cells compared to reconstituted cells. **Right:** Representation of each ribosomal protein color coded with low detection (red), normal detection (white) and high detection (blue). **(B)** Western blot analysis of whole cell lysate and isolated ribosomes in all cells showing normal level of ribosomal proteins in lysate, but decreased level of RPS15 and increased level of RPL5 in isolated ribosomes of TTD cells. More western blot analysis of more ribosomal proteins and quantification are given in the supplement Figure S3D-E. **(C)** Isolated ribosomes of WT, reconstituted and TTD cells were treated with 2M urea for 2 hrs. Exposed hydrophobic chains of unfolded proteins were quantified by Bis-ANS fluorescence. TTD cells show a higher amount of unfolded ribosomal proteins after partial denaturation. **(D)** Luciferase-based translation fidelity assay includes a control reporter Renilla and three Firefly reporters containing either no mutation (PC), with a variation for permanent inactivation (NC), and with a point mutatation K529N inactivating the Firefly (MUT). With correct translation, the point mutation is translated and Firefly is inactive, whereas with inaccurate translation the luminescence of the active Firefly is detected. The Firefly luminescence was normalized to the Renilla luminescence. **(E)** Plasmids of Renilla with either Firely NC or Firefly MUT were co-transfected via electroporation to all cells. Translation fidelity assay with plasmids shows increased translational infidelity in TTD cells. **(F)** Plasmids containing Renilla and either Firefly PC or Firefly MUT were transcribed by T7 polymerase and mRNAs were transfected with lipofectamine to all cells. Translation fidelity assay with mRNAs indicate increased translational infidelity in TTD cells. Data are represented as mean ± SD. ns p > 0.05, * p ≤ 0.05.

Sucrose gradient centrifugation was employed to isolate ribosomes to assess the unfolding stability of ribosomal proteins. The purified ribosomes were incubated with the chaotropic reagent urea and the subsequently exposed hydrophobic residues of unfolded ribosomal proteins were quantified by the fluorophore BisANS. The stability of proteins against unfolding by urea is a hallmark of long-lived species (Perez, Buffenstein et al., 2009, Treaster, Ridgway et al., 2014) and is considered as an indicator for translational accuracy (Azpurua, Ke et al., 2013). After urea challenge, ribosomal preparations of both TTD cell lines depict increased amounts of unfolded ribosomal proteins (Figure 3C) indicating a qualitative disturbance of the corresponding ribosomes. These results indicate that the amount of misfolded proteins in ribosomal preparations of TFIIEβ mutant cells is higher than in reconstituted and control cells.

To further unravel the functional base of the elevated load of misfolded proteins in ribosomal proteins, we assessed the error-rate of the translation process itself. For this purpose, a luciferase-based translation fidelity assay was used. This assay includes a control reporter *Renilla* and a *Firefly* reporter containing a point mutation K529N, which inactivates the *Firefly* luciferase (Figure 3D). The luciferase *Firefly* MUT stays inactive with correct, translation of the point mutation, whereas luminescence of the *Firefly* MUT is detectable only in the case that erroneous translation occurs. Plasmids expressing these constructs were co-transfected in WT, reconstituted and TTD cells (Figure 3E). Strikingly, both TTD cell lines show an increased luminescence indicative of a higher error rate when compared to the WT and reconstituted cells.

To exclude, that the re-activation of luciferase is due to errors in RNA polymerase II transcription, we transfected mRNA into cells. To further guarantee, that *Renilla*: *Firefly* luciferase are transfected in a ratio 1:1, plasmids expressing a fusion-protein of *Renilla* and *Firefly* were transcribed by means of a T7 kit and transfected to WT, reconstituted and TTD cells (Figure 3F). As observed in the translational fidelity assay with plasmid transfection, TTD cells depict similar high luciferase activity of *Firefly* MUT after mRNA transfection. We therefore conclude, that the TFIIEβ mutation in TTD cells fundamentally affects the accuracy of protein synthesis at the ribosome leading to the enrichment of misfolded proteins in the ribosome itself.

### TFIIEβ mutation result in a disturbed protein homeostasis and cell growth

Protein homeostasis is sustained by the balance of protein synthesis, maintenance and degradation. As our results unraveled a disturbance in ribosomal accuracy, we investigated the impact of translation on protein homeostasis in TTD cells. First we detected the rate of translational initiation by analyzing O-propargylpuromycin (OPP) incorporation in the growing aminoacid chain, which leads to a premature stop of translation. Interestingly, OPP-incorporation revealed an increased translational initiation in TTD cells (Figure 4A). Strikingly, the overall protein synthesis, which was assessed by metabolic labeling, was found to be significantly reduced in TTD cells. (Figure 4B). These data suggest that increased ribosomal inaccuracy leads to higher but ineffective protein synthesis initiation. This notion was further supported by the finding, that the cytoplasmic proteome of TTD cells is prone to heat denaturation. As shown in Figure 4C, more proteins are pelleted during centrifugation after heat-treatment, indicating an elevated load of misfolded proteins in TTD. Protein degradation analysis by measuring 20S proteasome activity revealed a markedly and significant reduction of the 20S proteasome activity in TTD cells (Figure 4D) that is in line with a reduced overall protein synthesis. Finally, we speculate that a disturbed protein homeostasis might also impact on cellular proliferation and compared growth properties of TTD cells with that of WT and reconstituted cells (Figure 4E). In comparison to WT cells, proliferation kinetics of TTD cells clearly revealed a reduced division rate in TTD cells when WT cells are in the exponential proliferation phase. Taken together, our data describe a severe disturbance in proteostasis as a novel pathomechanism which explains the observed reduced growth and proliferation rates in TTD cells, and which is most likely also responsible for the symptoms and clinical picture in TTD patients.

**Figure 4.**
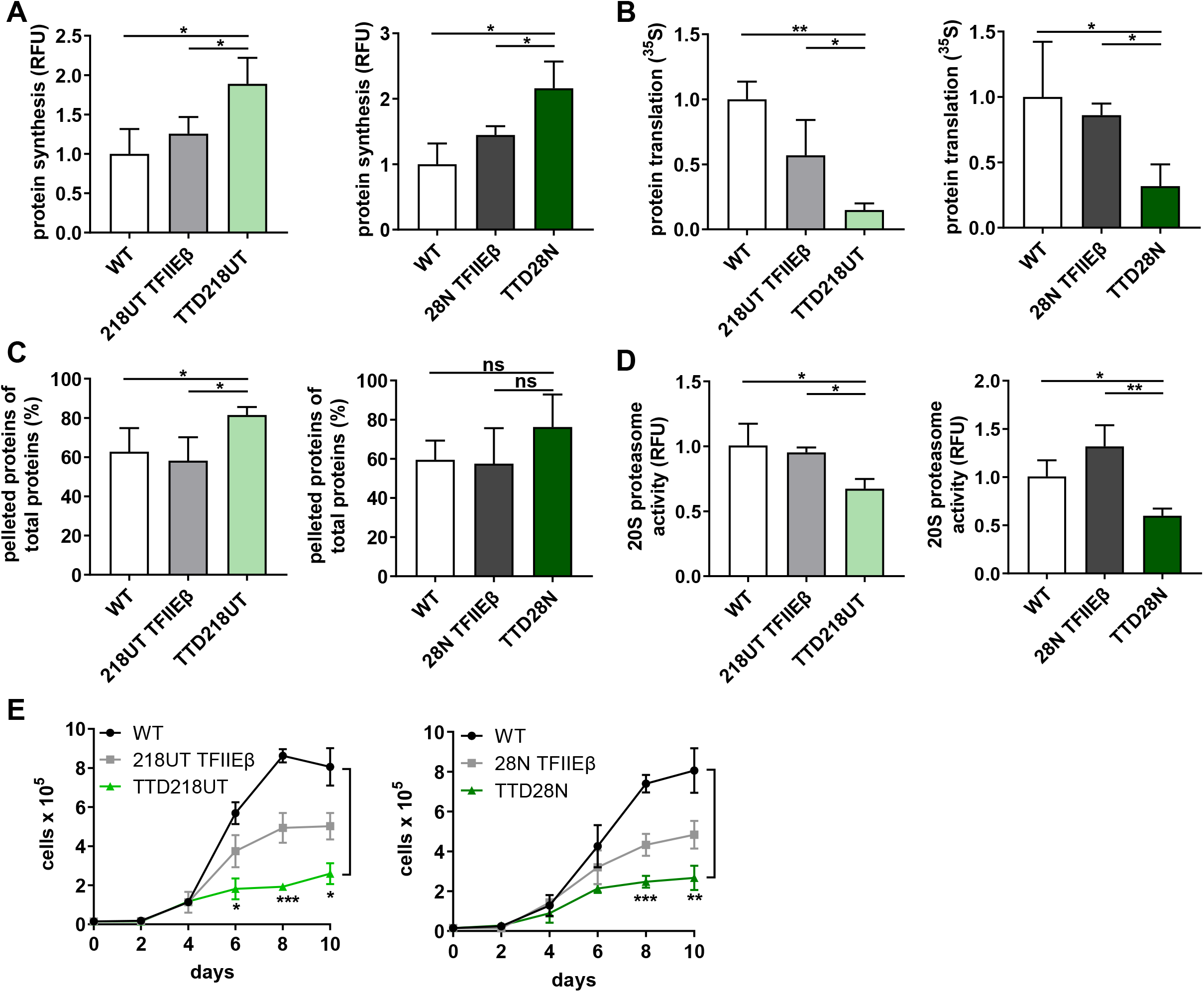
Loss of protein homeostasis in TTD cells. **(A)** 5 FAM-Azide detection of OPP-labeled proteins indicates the rate of protein synthesis initiation in WT, reconstituted and TTD cells. TTD cells show increased translation initiation. **(B)** Analysis of total protein translation by using ^35^S metabolic labeling shows decreased protein synthesis in TTD cells compared to WT and reconstituted cells. **(C)** Heat sensitivity analysis of cytoplasmic extract in WT, reconstituted and TTD cells shows increased percentage of pelleted proteins in TTD cells after heat treatment for 15 min at 99 °C. **(D)** Detection of cleaved 20S specific substrate, SUC-LLVY-AMC, indicating the 20S proteasome activity in WT, reconstituted and TTD cells. Reduced proteasome activity was observed in TTD cells. **(E)** Proliferation kinetics of TTD cells compared to WT and reconstituted cells indicates reduced cell proliferation in TTD cells. Data are represented as mean ± SD. ns p > 0.05, * p ≤ 0.05, ** p ≤ 0.01, *** p ≤ 0.001.

## Discussion

The heterodimeric general transcription factor TFIIE recruits TFIIH to the preinitiation complex of RNA polymerase II promoters and is thus involved in gene expression of all protein-coding genes (Compe et al., 2019, Ohkuma, Sumimoto et al., 1990). Here we describe an unanticipated role of TFIIE which profoundly impairs ribosomal biogenesis. Our data shows enrichment of TFIIE in the nucleolus, at the site of RNA polymerase I transcription. TFIIE enrichment depends on RNA polymerase I activity, as it’s inhibition by CX5461, clearly reduces nucleolar TFIIE staining. So far, a nucleolar localization and function of general transcription factors of RNA polymerase II was reported for TFIIH (Hoogstraten, Nigg et al., 2002, Iben et al., 2002). A seminal publication indicates that nucleolar RNA polymerase II transcription is essential for proper rRNA biogenesis and processing (Abraham, Khosraviani et al., 2020). The authors showed that RNA polymerase II transcribes intergenic spacers (IGSs) of the rDNA to maintain nucleolar organization and hence rRNA synthesis and maturation. However, it is largely unresolved how RNA polymerase II transcribes at the IGSs of the rDNA, and which general factors are involved. We here addressed the unique possibility that the nucleolar TFIIE enrichment is due to nucleolar RNA polymerase I transcription. In fact, our data unveil that the coding region of the rDNA, that is actively transcribed by RNA polymerase I, is bound by TFIIE and selective inhibition of RNA polymerase I transcription reduces TFIIE binding. These data highlight a direct involvement of TFIIE in RNA polymerase I transcription. The C-terminal end of TFIIEβ is capable of binding to DNA (Okamoto, Yamamoto et al., 1998), suggesting that TFIIE may be bound to the rDNA through its small subunit TFIIEβ. The fact that the mutation of TFIIEβ weakens the rDNA gene-occupancy of RNA polymerase I, indicates that TFIIE has a direct impact on the process of ribosomal biogenesis. Further investigations will reveal, if reduced binding of the elongation factor TFIIH to the rDNA is also observed, as TFIIE recruits TFIIH in RNA polymerase II pre-initiation complex formation (Roeder, 1996). A reduced gene-occupancy of RNA polymerase I, as shown in this study, was already described by Nonnekens et al (Nonnekens et al., 2013) for TTD and CS-causing mutations in TFIIH and CSB indicating that this might be a common pathophysiology in these childhood syndromes. Moreover, in this publication the authors further showed, that co-transcriptional processing of the pre-rRNA is disturbed in cells and tissues from a TTD mouse model with a mutation in TFIIH. The process of RNA polymerase I transcription elongation is coupled to processing and pre-ribosomal assembly (Bohnsack & Bohnsack, 2019, Scull & Schneider, 2019), thus disturbances in transcription elongation might impact on the subsequent maturation steps. In our study we demonstrate a previously unreported accumulation of processing intermediates and a reduced abundance of the 18S mature rRNA in TFIIEβ cells. Maturation disturbances as a consequence of transcription elongation defects in the rRNA can also translate to ribosomal assembly defects as described in yeast (Schneider, Michel et al., 2007). We compared ribosomal composition of patient cells with reconstituted cells and detected an underrepresentation of ribosomal proteins of the small ribosomal subunit. The small ribosomal subunit 40S is the translation initiating and codon-decoding subunit of the ribosome. Disturbed ribosomal composition can lead to decoding-defects of the ribosome (Lezzerini, Penzo et al., 2020, Paolini, Attwood et al., 2017, Venturi & Montanaro, 2020) detectable by luciferase-based translation assays. We took advantage of a well-established luciferase transfection system (Azpurua et al., 2013) and discovered an elevated error-rate of the translation process in TTD patient cells. The reduced translational fidelity, described for the first time in TTD, has recently been reported in CS-patient derived fibroblasts (Alupei et al., 2018). We here found translational infidelity followed by a loss of proteostasis and disturbed protein synthesis in TTD, indicating that TTD and CS share distinct aspects in their pathophysiology and this may be a more general mechanism in both childhood syndromes. In this regard, both are caused by mutations in the same subunits of the transcription/DNA repair factor TFIIH. Additionally, these similarities might suggest, that the loss of proteostasis in CS might not origin from DNA-damage, as a comparable defect is here shown for TTD cells which do not have any deficiencies in Nucleotide Excision Repair (Theil et al., 2017).

The loss of proteostasis is also evident by the elevation of heat-sensitive proteins in patient cells. Heat-stability of the proteome is a hallmark of long-living species and characterizes the longest-living animal, *Arctica Islandica* (Treaster et al., 2014). Elevated heat sensitivity indicates that a detectable proportion of the proteome is affected by misfolding, and this most likely is caused by translational errors. In support of this, ribosomal proteins display an elevated unfolding by urea in comparison to control ribosomes, indicating that the ribosomes themselves are affected by misfolded proteins likely due to an error-prone translation process. Our findings re-vitalize the error catastrophe theory of aging (Orgel, 1963). In his theory, Orgel postulates that elevated translational infidelity perpetuates and enhances itself by the accumulation of errors in the amino acid sequence of ribosomal proteins that have influence on translational fidelity.

The reduced translational fidelity at the ribosome in TFIIEβ mutant cells might be the cause of the observed loss of proteostasis. Interestingly, we detected an increase in the initiation of protein synthesis in TTD cells, while total protein synthesis is significantly reduced. This supports the assumption that the translation process is inefficient and error-prone in TFIIEβ cells. TTD is mainly regarded as a transcriptional syndrome, as 50% of the cases do not display DNA-repair disturbances (Stefanini, Botta et al., 2010). We here extend this classification, and highlight a possible contribution of protein translation disturbances in the pathophysiology of TTD. Recently, mutations in aminoacyl-tRNA synthetases were identified to cause TTD (Kuo et al., 2019, Theil et al., 2019) implying that a disturbed translation process, as shown in this study, might be causal for the developmental delay and neurodegeneration observed in TTD. This assumption is based on the hypothesis that only disturbances in a common cellular signaling pathway that is affected by all genes in this genotypical heterogenous disease can explain the phenotype. Here we provide first evidence, that the translation process at the ribosomes might be this common pathway. If also mutations in the TTD-causing genes TTDN1 (MPLKIP) and RNF113A are impacting on the translation process awaits further analysis.

## Acknowledgements

We thank Arjan Theil and Wim Vermeulen from the Department of Molecular Genetics, Oncode Institute, Erasmus University Medical Center, University Medical Center Rotterdam (Netherlands) for providing the cell line TTD218UT and the plasmid pLenti_puro_CMV_TFIIEβ-GFP; Donata Orioli from the Institute of Molecular Genetics, National Research Council Pavia (Italy) for providing the cell line TTD28N; Markus Schosserer from the Institute for Molecular Biotechnology, University of Natural Resources and Life Sciences Vienna (Austria) and Vera Gorbunova from the Department of Biology, University of Rochester (USA, NY) for providing Firefly and Renilla plasmid constructs for the luciferase-based translation fidelity assay. We used the confocal microscope and ultracentrifuge in the Institute of Molecular Genetics and Cell Biology, University Ulm (Germany) and thank Nils Johnsson, Thomas Gronemeyer and Matthias Hecht for permission and support. We are grateful Harsh Ranawat from the Institute for Biophysics, University Ulm (Germany) for providing antibodies and reagents for immunofluorescence, taking high resolution STED images and scientific discussion. We further thank Lisa Wiesmüller from the Department of Gynecological Oncology, University Hospital Ulm (Germany), Ambra Giglia-Mari from the Institute for Neuromyogene, University Claude Bernhard Lyon 1 (France), Meinhard Wlaschek for scientific discussion and input. Sebastian Iben was supported by a grant of the German Research Foundation DFG (IB83/3-4), and Tamara Phan by the DFG program CEMMA (cellular and molecular mechanisms of aging GRK1789).

## Author Contributions

Conceptualization, S.I.; Methodology, S.I., T.P., and P.M.; Investigation, T.P., L.S., and C.L.; Writing – Original Draft, T.P., and S.I.; Writing – Review and Editing, S.I.; Visualization, T.P.; Funding Acquisition, S.I. and K.S.-K.; Resources, S.I., J.M., and K.S.-K.; Supervision; S.I., P.M., J.M., K.S.-K.

## Materials Availability

All unique/stable reagents generated in this study are available from the Lead Contact with a completed Materials Transfer Agreement.

## Methods

### Reagents and resources

Antibodies and oligonucleotides are given in **Error! Reference source not found.**-Table S 2.

### Cell culture

Healthy WT controls were grown in DMEM (41965-039, Gibco) supplemented with 10% FBS and 1% penicillin-streptomycin. SV40-immortalized human NP-TTD fibroblasts (TTD218UT, TTD28N) were cultured in DMEM/HAM-F10 (P04-12500, PAN BioTech) in a ratio 1:1 supplemented with 10% FSB and 1% penicillin-streptomycin. Reconstituted cells (218UT TFIIEβ, 28N TFIIEβ) were grown in DMEM/HAM-F10 (1:1) supplemented with 10% FBS, 1% penicillin-streptomycin and 2 µg/ml puromycine. All cells were cultured at 37 °C and 5% CO_2_. For RNA polymerase I transcription inhibition, cells on culture dishes with 80 % cell density were incubated with 1µM CX5461 for 4 hrs.

### Sequence analysis of D187Y variation

The homozygous point mutation in NP-TTD genome was verified by primers binding 100 nucleotide up- and downstream from the variation (forward: 5’ GTATACTGAGTGTGTTAAGGAAATG 3’; reverse 5’ AAAGCTCACCTTCATCCAC 3’). PCR product was sent for sequencing and data were analyzed by using CLC Workbench 7.7.3.

### Reconstitution of 218UT TFIIEβ and 28N TFIIEβ cell lines

NP-TTD cells were reconstituted with a plasmid overexpressing wild-type TFIIEβ-GFP (pLenti_puro_CMV_TFIIEβ-GFP). 2×10^6^ NP-TTD were transfected with 3 µg plasmid via electroporation using Neon™ Transfection System (MPK1096, Invitrogen) with following parameters: 1100V, 20ms and 2x pulses. Plasmid transfection was verified by flow cytometry. Cells were washed in PBS and flow cytometry analysis were performed in FACS buffer (PBS including 2% FBS). The data were analyzed with FlowJo Software.

### Immunofluorescence

Cells were seeded on pyrolysine coverd slides for 1 day. 80% confluent cells were washed with PBS, fixed with 4% paraformaldehyde (4°C) for 15 min, washed with PBS, permeabilized with 0.3 % Triton X-100/PBS and blocked at RT for 1 hr with 5% BSA including 10% goat serum. Antibodies were diluted in Dako Antibody Diluent and cells were incubated with primary antibodies at 4 °C overnight in a moist chamber. Cells were washed with PBS, incubated with secondary antibodies at RT for 1 hr in a moist chamber, washed with PBS and incubated at RT for 1hr with the DNA probe SPY650-DNA (SC501, SpiroChrome). After washing with PBS, cells were embedded in Dako mounting medium. Images were taken by fluorescence microscope (Zeiss Axio Imager M1, 64 oil objective), confocal microscope (Zeiss Axio Observer Z1 confocal laser scanning microscope, 100 oil objective) and processed and quantified by using ImageJ.

### Super-resolution microscopy

Super-resolution optical microscopy images were taken using a custom built dual-color 3D STED microscope (Osseforth, Moffitt et al., 2014). Therefore, 3 × 10^4^ cells were seeded on glass Ibidi µ-slide 8 well glass-bottom plates and incubated o/N. Cells were subsequently fixed with EM-grade PFA and immunostaining was performed as described above. Finally, cells were mounted in 97% 2,2’-thiodiethanol (TDE) for index matching. Antibodies used for immunofluorescence are given in Table S 1. Typically, an excitation power of 1 µW and a depletion power of 1.3 mW and a repetition rate of 1Mhz were used. The STED images have pixel size of 20 nm and were captured with a pixel dwell, time of 300 µs. Images were analyzed using ImageJ 1.51f and a Gaussian Blur of σ = 1 was applied. For better visualization brightness and contrast settings were adapted for the images.

### Real-time qPCR standard curve analysis

1 µg of total RNAs were reverse transcribed with Moloney murine leukemia virus Reverse Transcriptase (M170B, Promega). 100 ng cDNA and FastStart Universal SYBR Green Master (04913850001, Roche Diagnostics) were used for real-time qPCR analysis (denaturation at 95 °C for 10s, annealing at 60 °C – 68°C for 30s, elongation at 72 °C for 30s). A prepared standard curve of the oligonucleotide of interest with a linear regression with R^2^-values >0.8 was used for calculation of the absolute amount (ng) of the oligonucleotide of interest within 100ng total cDNA. Primers used for qPCR analysis are listed in Table S 2.

### Chromatin Immunoprecipitation (ChIP)

80% confluent cells were fixed with 1% formaldehyde (28908, Thermofisher Scientific) at RT for 10 min and 0.125 M glycine were used to stop cross-linking reaction. Cells were washed with PBS, harvest and lysed with ChIP cell lysis buffer (5mM HEPES pH 8.0, 85 mM KCl, 0.5 % NP-40 (Pierce), and 1:50 cOmplete proteinase inhibitor cocktail mix (Roche)) on ice for 10 min. Pelleted Chromatin was sonicated in ChIP sonicate buffer (50 mM HEPES pH 7.9, 140 M NaCl, 1 mM EDTA, 1 % Triton X 100, 0.1 % Na-deoxychlorate, 0.1 % SDS, 0.5 mM PMSF, 1:50 cOmplete) by a Focused-ultrasonicator (Covaris M220, 1 ml) to yield chromatin fragments with an average size of 600 bp.

For chromatin precipitation, Protein A agarose beads (20334, Pierce) or Protein A magnetic dynabeads (10002D, Invitrogen) were used. After incubating 10 µg chromatin with antibodies at 4 °C overnight in IP diluent (0.1% SDS, 1 % Triton X 100, 1 mM EDTA pH 8.0, 16.7 mM Tris-Cl pH 8.0, 167 mM NaCl, 1:50 cOmplete), chromatin-antibody-complexes were washed first with low salt buffer (0.1 % SDS, 1 % Triton X 100, 2 mM EDTA, 20 mM Tris-Cl, pH 8.1, 150 mM NaCl), then with high-salt buffer (0.1 % SDS, 1 % Triton X 100, 2 mM EDTA, 20 mM Tris-Cl, pH 8.1, 500 mM NaCl), with LiCl buffer (10 mM Tris-Cl pH 8.0, 250 mM LiCl, 1% NP40, 1% deoxychloric acid, 1 mM EDTA). and twice with TE buffer (10mM Tris pH 8, 1mM EDTA). ChIPed DNA was eluted from beads in µChIP elution Buffer (10 mM Tris-Cl pH 8, 1 mM EDTA pH 8, 1 % SDS, 1 % proteinase K) for 2 hrs at 60 °C with 900 rpm. ChIPed DNA was purified by using QIAquick® Nucleotide Removal Kit (28306, Qiagen) according to the manufactor’s protocol. Samples were qualitative or quantitative analyzed via PCR or qPCR, respectively. For qualitative ChIP analysis, samples were loaded on 1% agarose gel and visualized by using Ged-Doc-It Imager (UVP). For sequential ChIP analysis (Re-ChIP), samples were released after the first round of precipitation by incubation in 10 mM DTT for 30 min at 37 °C at 500 rpm and incubated with the respective other antibody for the second round of precipitation. Primers and antibodies used for ChIP analysis are given in Table S 1 - Table S 2.

### Western Blot analysis

Cells were grown on 15 cm culture dishes until 80 % cell density, harvest and lysed with 100 µl lysis buffer (10% glycine, 1 % Triton X 100, 137 mM NaCl, 20 mM Tris pH 8, 2 mM EDTA pH 8.1. 100 – 300 µg protein was loaded on 10 – 12% SDS-PAGE and transferred at 4°C overnight to a nitrocellulose blotting membrane (A29434119, GE Healthcare) in transfer buffer (25 mM Tris pH 8, 192 mM glycine, 5% methanol). Membranes were blocked at RT for 1 hr with 5 % milk powder and 0.1 % Tween 20 (diluted in PBS), washed in PBS, incubated with primary antibodies at 4 °C overnight, washed with, PBS and incubated with secondary antibodies at RT for 1 hr. Membranes were developed using Fusion Fx7 (Vilber). Images were processed and quantified by using ImageJ. Anitbodies used for western blot analysis are listed in Table S 1.

### Methylation sensitivity restriction analysis

Isolated Chromatin was digested with 1% proteinase K for 2 hrs at 60 °C with 300 rpm and was subsequently purified by using QIAquick® Nucleotide Removal Kit (28306, Qiagen) according to the manufactory’s protocol. 1 µg DNA was digested with either the methylation-sensitive restriction enzyme HpaII (1x Buffer A, 1x BSA) or the methylation-insensitive restriction enzyme MspI (1x Buffer B, 1x BSA) for 1 hr at 37 °C. For qualitative analysis via PCR (denaturation at 95 °C for 30s, annealing at 58°C for 30s, elongation at 72 °C for 30s), primers amplifying the promoter region of rDNA were used and loaded on 1 % agarose gel. Gel was detected with GelDoc-It Imager (UVP).

### Northern Blot analysis

5 µg total RNA were denaturated at 65 °C for 15 min in RNA loading dye (50% formamide, 7.5 % formaldehyde, 1x MOPS, 0.5% ethidiumbromide) and immediately placed on ice for 5 min. Samples were separated on a 0.9 % agarose gel (diluted in 1x MOPS) with 80V for 2.5-3 hrs. The gel was soak in 20x SSC (3M NaCl, 0.3 M Na_3_C_6_H_5_O_7_, adjust to pH 7) and RNAs were transferred to Amersham Hybond membrane (RPN303S, GE Healthcare) soaked in 2x SSC. Membrane was pre-hybridized for 2 hrs at 60°C with pre-hybridization buffer (50 % formamide, 0.1 % SDS, 8x Denhards solution, 5x SSC buffer, 50 mM NaP buffer, 0.5 mg/ml t-RNA). Probes were synthesize by T4 polynucleotide kinase using radioactive [^32^P] γATP. Membranes were hybridize with radioactive ^32^P-labeled oligonucleotides using pre-hybridization buffer at 60 °C for 1 hr and subsequently at 37 °C overnight. Finally, membranes were exposed to X-ray film and quantified with ImageJ after Wang and Pestov (**Error! Reference source not found.**). For rRNA processing pathway analysis we used probes binding to the region ITS1 (5’ GGGCCTCGCCCTCCGGGCTCCGTTAATGATC 3’) and ITS2 on the rDNA (5’ CTGCGAGGGAACCCCCAGCCGCGCA 3’).

### Ribosome preparation

Ribosome preparation was performed after Penzo et. al. (**Error! Reference source not found.**). 80% confluent cells were washed, harvest and lysed with cell lysis buffer (10mM Tris-HCl pH 7.5, 10 mM NaCl, 3mM MgCl_2_, 0.5 % Nonidet P40) on ice for 10 min. Samples were centrifuged for 12 min 16900 xg and lysates were incubated with protein synthesis master mix (150 mM HEPES KOH pH 7.5, 400 mM Kcl, 250 µM L-amino acid mixture, 1.25 mM GTP, 10 mM ATP, 25 mM creatine phosphate, 2mM spermidine, 2.5 mM DTT, 5mM magnesium acetate, 0.9 mg/ml creatine kinase) for 10 min at 37 °C. Subsequently, samples were transferred to a sucrose gradient, consisting of a bottom sucrose layer (30 mM HEPES KOH pH 7.5, 70 mM KCl, 2 mM magnesium acetate, 1 mM DTT, 1M sucrose) and a high-salt-top layer (30 mM HEPES KOH pH 7.5, 0.5 M KCl, 2mM magnesium acetate, 1mM DTT, 0.7 M sucrose). After ultracentrifugation (Optima™ MAX-XP Ultracentrifuge, Beckman Coulter) for 15 hrs at 4 °C with 110 000 xg, pelleted ribosomes were resuspended in ribosome solution (10 mM Tris HCl pH 7.5, 2mM magnesium acetate, 100 mM ammonium acetate).

### Ribosome stability assay using BisANS

5 µg isolated ribosomes were treated with 2M of the detergent Urea for 2 hrs. Conformational changes of proteins were detected by the probe BisANS (4,4’-Dianilino-1,1’-Binaphthyl-5,5’-Disulfonic Acid, Dipotassium Salt). BisANS is nonfluorescent in water and only becomes appreciably fluorescent when bound to the hydrophobic site of proteins. Hence, BisANS is a sensitive indicator of protein unfolding. Fluorescent was detected by a plate reader Varioskan™ LUX (Thermofisher Scientific) using excitation 375 nm / emission 500 nm.

### Mass spectrometry analysis

Mass spectrometry analysis of isolated ribosome from NP-TTD cell line TTD218UT and reconstituted cell line 218UT TFIIEβ were performed by the Bavarian Center for Biomolecular Mass Spectrometry, Technical University Munich. Label-free quantification of mass spectrometry data were analyzed using Excel and the program software R Studio.

### Luciferase-based translation fidelity assay with plasmid transfection

The translation fidelity assay via plasmid transfection includes the control reporter luciferase *Renilla* and experimental reporters *Firefly* (Neg, Mut). *Renilla* and *Firefly* were <u>expressed on separate plasmids</u> (kindly provided by Vera Gorbunova). Cells were cultured in 10 cm culture dishes until 80 % cell density. 15 x10^4^ cells were harvest and co-transfected with 0.1 µg control plasmid *Renilla* and 5µg reporter plasmid *Firefly* (Neg or Mut) via electroporation using Neon™ Transfection System (MPK1096, Invitrogen) with following parameters: 1100V, 20ms and 2x pulses. Subsequently, cells were seeded in 5×10^4^ cells/well in white 96 well plates and were grown 24 hrs. Luciferase activities were detected by using Dual-Glo® Luciferase Assay System (E2920, Promega) according to the manufactory’s protocol.

### Luciferase-based translation fidelity assay with mRNA transfection

The translation fidelity assay via mRNA transfection includes the control reporter luciferase *Renilla* and experimental reporters *Firefly* (Pos or Mut). *Renilla* and *Firefly* were <u>expressed on one plasmid</u> (kindly provided by Markus Schosserer). Plasmids were transcribed to mRNAs by using Ampli Cap-Max T7 High Yield Messager Marker Kit (C-ACM04037, CellScript) according to the manufactory’s protocol. 10^5^ cells/well in 100 µl culture medium were seeded in white 96 well plate and were grown overnight. 500ng mRNA/well in 50 µl OptiMEM (31985070, Gibco) and 1 µl/well of Lipofectamine® MessengerMAX mRNA Transfection Reagent (LMRNA003, Invitrogen) in 50µl OptiMEM were incubated for 10 min at RT. mRNA dilution and Lipofectamin dilution were mixed and incubated for further 15 min at RT. After removing the old media from cells, 100 µl of mRNA-Lipofectamin-OptiMEM mixture were transferred to each well and cells were grown for 24 hrs. Luciferase activities were detected by using Dual-Glo® Luciferase Assay System (E2920, Promega) according to the manufactory’s protocol.

### Protein synthesis analysis

Protein synthesis was detected by using Protein Synthesis Assay Kit (601100, CaymanChemical) according to the manufactory’s protocol. Briefly, 2.5 x10^4^ cells/ml were grown in white 96 well plate overnight, incubated with OPP (O-Propargyl-puromycin) working solution for 1 hr, fixed, and stained with 5 FAM-Azide solution for 30 min. OPP binds to newly synthesized polypeptides and 5 FAM-azide is fluorescent upon binding to OPP. Fluorescence was detected by using a plate reader Varioskan™ LUX (Thermofisher Scientific) using excitation 485 nm / emission 535 nm.

### ^35^S Methionine Labeling

Protein translation was analyzed by using metabolic labeling of cells with ^35^S methionine. Therefore, 5 x10^4^ cells per reaction were harvest in 1.5 ml reaction tube by centrifuging for 5 min at 3000 rpm, and incubated at 37 °C for 30 min in 100 µl methionine-free DMEM (21013-024, Gibco) supplemented with 10% FBS and 1% penicillin-streptomycin. Subsequently, 10 µl culture medium with 1% ^35^S-labeled methionine was added to cells and cells were incubated for further 1 hr. After centrifuging cells for 5 min at 3000 rpm, cell pellet was resuspened in 100 µl 1mg/ml BSA and each reaction was vortex for 3 sec. 1 ml of cold 10% TCA was added, vortexed for 3 sec and reactions were incubated on ice for 1 hr. Reaction were transferred onto glass fibre prefilters (Merck, APFC02500) and washed with cold 10% TCA and subsequently with cold 70% ethanol. Glass fibre prefilters were air dried for 5 min and the amount of ^35^S-labeled methionine was analyzed with 5 ml liquid scintillation cocktail Ultima Gold™ (6013329, PerkinElmer) by Liquid Scintilation Analyzer Tri-Carb 4910 TR (PerkinElmer).

### 20S Proteasome activity analysis

20S Proteasome activity was determined by using 20S Proteasome Assay Kit (10008041, CaymanChemical) according to the manufactory’s protocol. In short, 5x 10^5^ cells/well in 100 µl culture medium were seeded in white 96 well plate, lysed, and incubated with a 20S specific substrate solution for 1 hr. The 20S specific substrate is fluorescent upon cleavage and fluorescence was detected by using a plate reader Varioskan™ LUX (Thermofisher Scientific) using excitation 360 nm / emission 480 nm.

### Heat Sensitivity analysis

Heat sensitivity analysis was adapted from Treaster and colleagues (**Error! Reference source not found.**). Cells were grown on 15 cm culture dished to 80 % cell density. 4×10^6^ cells were harvest and its PCV (packed cell volume) was estimated. Cells were dissolved in 1.5x PCV of Dignam A buffer (10 mM KCl, 10 mM Tris pH 7.9, 1.5 mM MgCl_2_, 1:50 cOmplete proteinase inhibitor mix) for 15 min on ice. Cells were lysed by passing cells 50 times through a 23G syringe. Cytoplasmic extract was separated from nuclei and cell debris by centrifugation for 5 min at 4000 rpm at 4 °C and subsequently by ultracentrifuge for 1 hr with 100,000 xg at 4 °C. 150µg of protein lysate was heat treated at 99ºC for 15 min and subsequently centrifuged for 5 min with 14 000 rpm at 4 °C. The pellet was resuspended in 10 µl 4M Urea and the protein concentration of both pellet and supernatant was determined by using the method of Bradford (5000006, Bio-Rad). Finally, the percentage of pelleted protein in relation to total protein was calculated.

### Growth kinetic

2 × 10^5^ cells were seeded per well in a 6 well culture plate. For each time point 3 wells were detached using 1x trypsin and manually counted using a haemocytometer (Neubauer chamber, 0.100 mm depth, 0.0025 mm^2^ area). Cells were counted every 48 hrs over a period of 10 days.

## Figure Supplement legends

**Figure S1.**
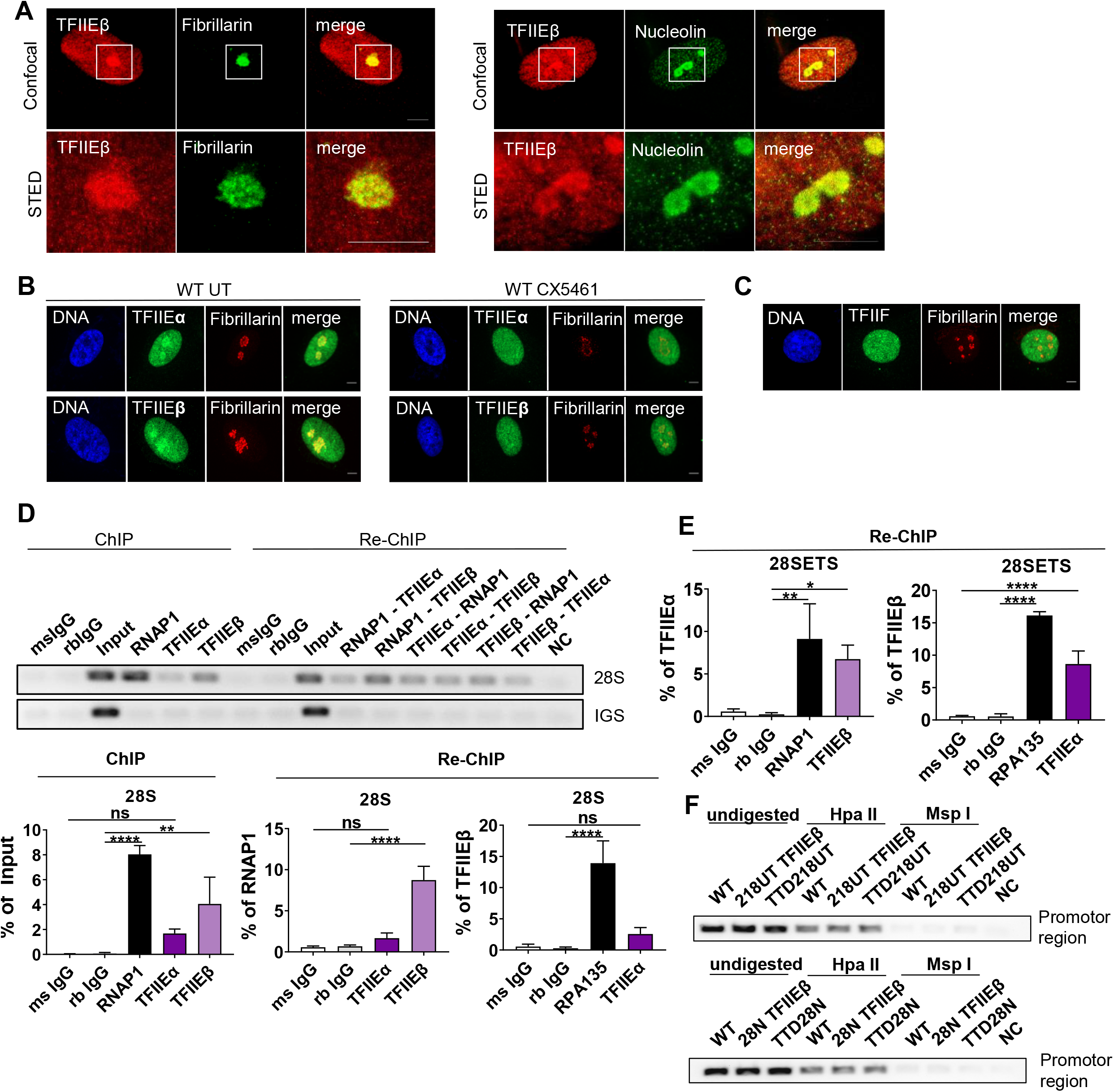
Enrichment of TFIIE in the nucleolus and binding to the rDNA. **(A)** High resolution STED microscopy images indicating co-localization of TFIIEβ with Fibrillarin (top) and Nucleolin (bottom). Scale bar 5 µm. **(B)** Confocal immunofluorescence microscopy (100X) of WT cells shows co-localization of both TFIIE subunits with Fibrillarin. Inhibition of RNA polymerase I transcription in WT cells by 1 µM CX5461 result in a re-distribution of both TFIIE subunits from the nucleolus. Scale bar 5 µm. **(C)** TFIIF staining in confocal immunofluorescence microscopy (100X) of WT cells shows no nucleolar enrichment. Scale bar 5 µm. **(D)** Qualitative (top) and quantitative (bottom) qPCR analysis of ChIPs indicate binding of TFIIEβ to the rDNA at the 28S coding region. Sequential ChIP analysis show binding of TFIIEβ to the same rDNA molecule as RNAP1 at the 28S coding region of the rDNA. **(E)** Quantitative qPCR analysis of Re-ChIP confirms the significant association of both TFIIE subunits and RNA polymerase I at the 28SETS region of the rDNA. **(F)** Methylation sensitive restriction analysis of genomic DNA shows wt-levels of active and inactive rDNA in TTD cells.

**Figure S2.**
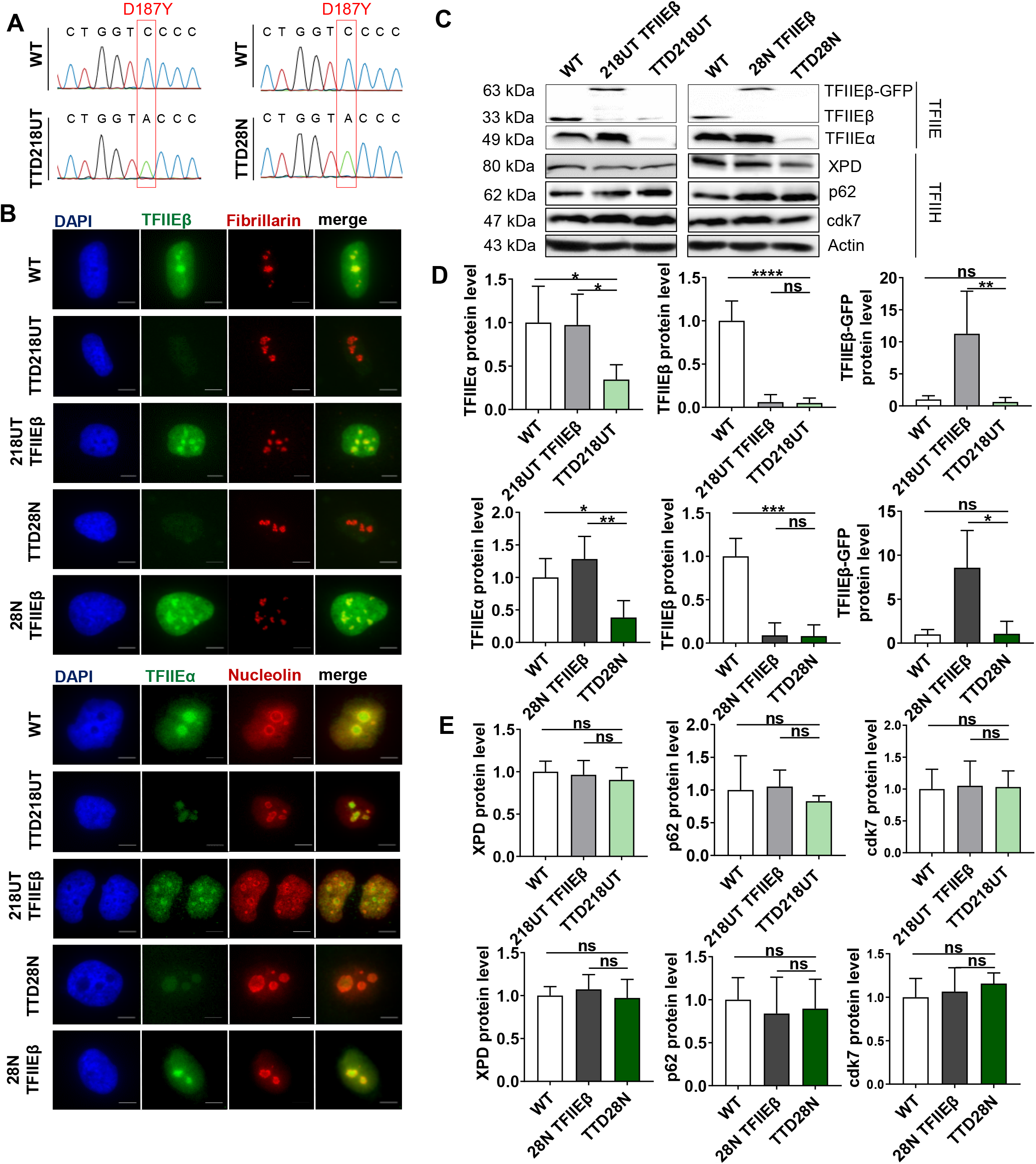
Reduced TFIIE protein level in TTD cell line. **(A)** Sequence analysis of TTD218UT (left) and TTD28N (right) compared to WT shows a homozygous mutation (c.559C>A [p.Asp187Tyr]) in both TTD cell lines. **(B)** Immunofluorescence microscopy (100X) of WT cells reveal co-localization of TFIIE and nucleolus marker including Fibrillarin and Nucleolin in WT and reconstituted cells. Reduced fluorescence signals of both TFIIE subunits were observed in TTD cells. **(C)** Western Blot analysis indicates reduced protein abundance of both TFIIE subunits in TTD cells compared to WT and reconstituted cells, whereas the protein abundance of TFIIH is unaffected in TFIIEβ-mutated cells. **(D-E)** Quantification of western blot analysis in Figure S2 C indicate significant reduced protein level of both TFIIE subunits and wild-type protein level of TFIIH.

**Figure S3.**
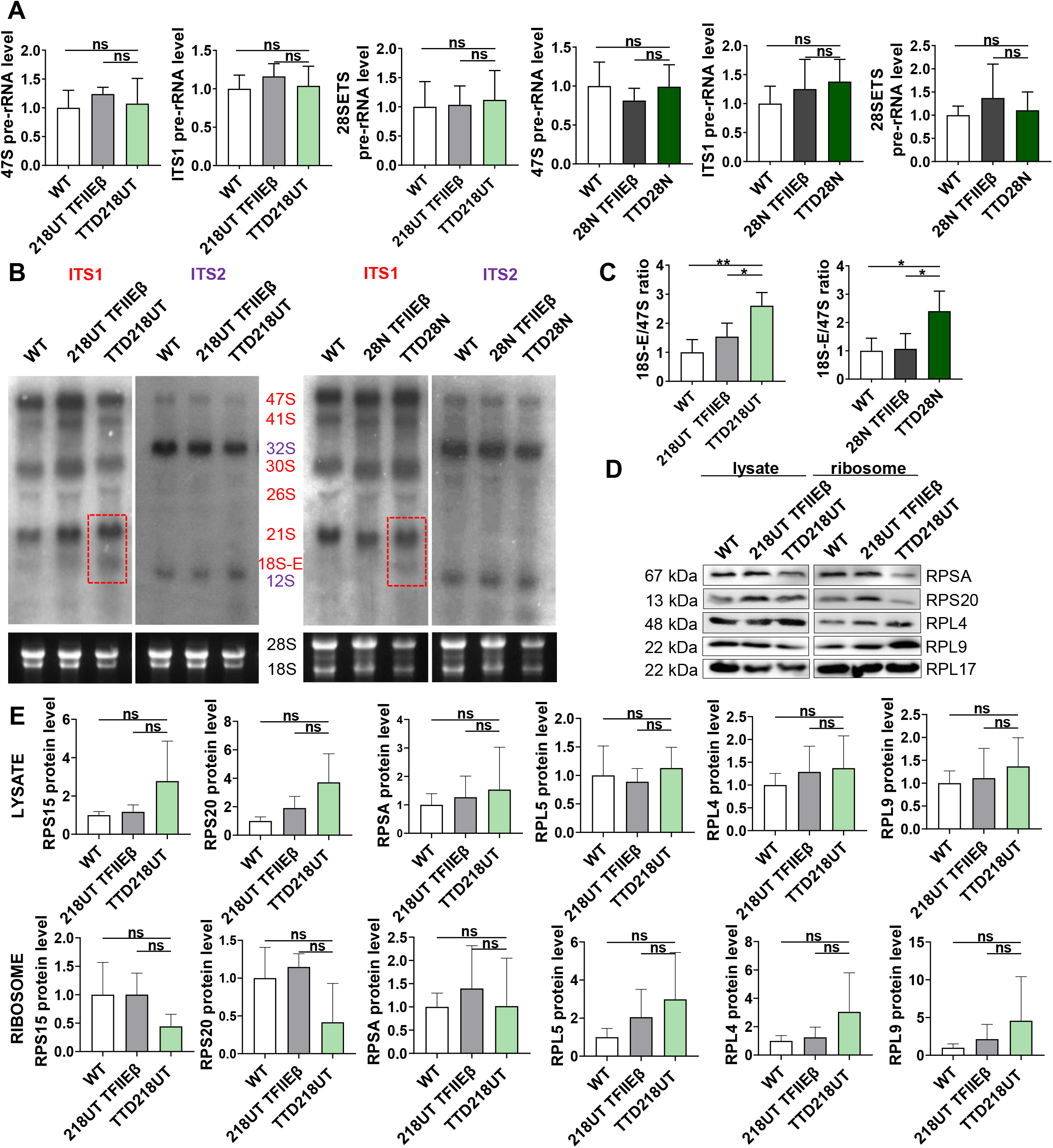
Disturbed rRNA processing and ribosomal composition in TTD cells. **(A)** Quantitative qPCR analysis of different regions of the RNA polymerase I primary transcript indicates wild-type levels of the pre-rRNA. **(B)** Full images of northern blot analysis. Membranes were probed with ITS1 (red), stripped, and re-probed with ITS2 (purple). **(C)** Quantification of 18S-E/47S ratio in WT, reconstituted and TTD cells indicating increased 18S-E level in TTD cells. **(D)** Western blot analysis of whole cell lysate and isolated ribosome in WT, reconstituted 218UT TFIIEβ cells and TTD218UT cells. **(E)** Quantification of Western Blots of three ribosomal preparations and selected ribosomal proteins point towards a tendency of over- and underrepresented proteins in lysates and ribosomal preparations.

## Supplemental material: Tables S

**Table S1.**
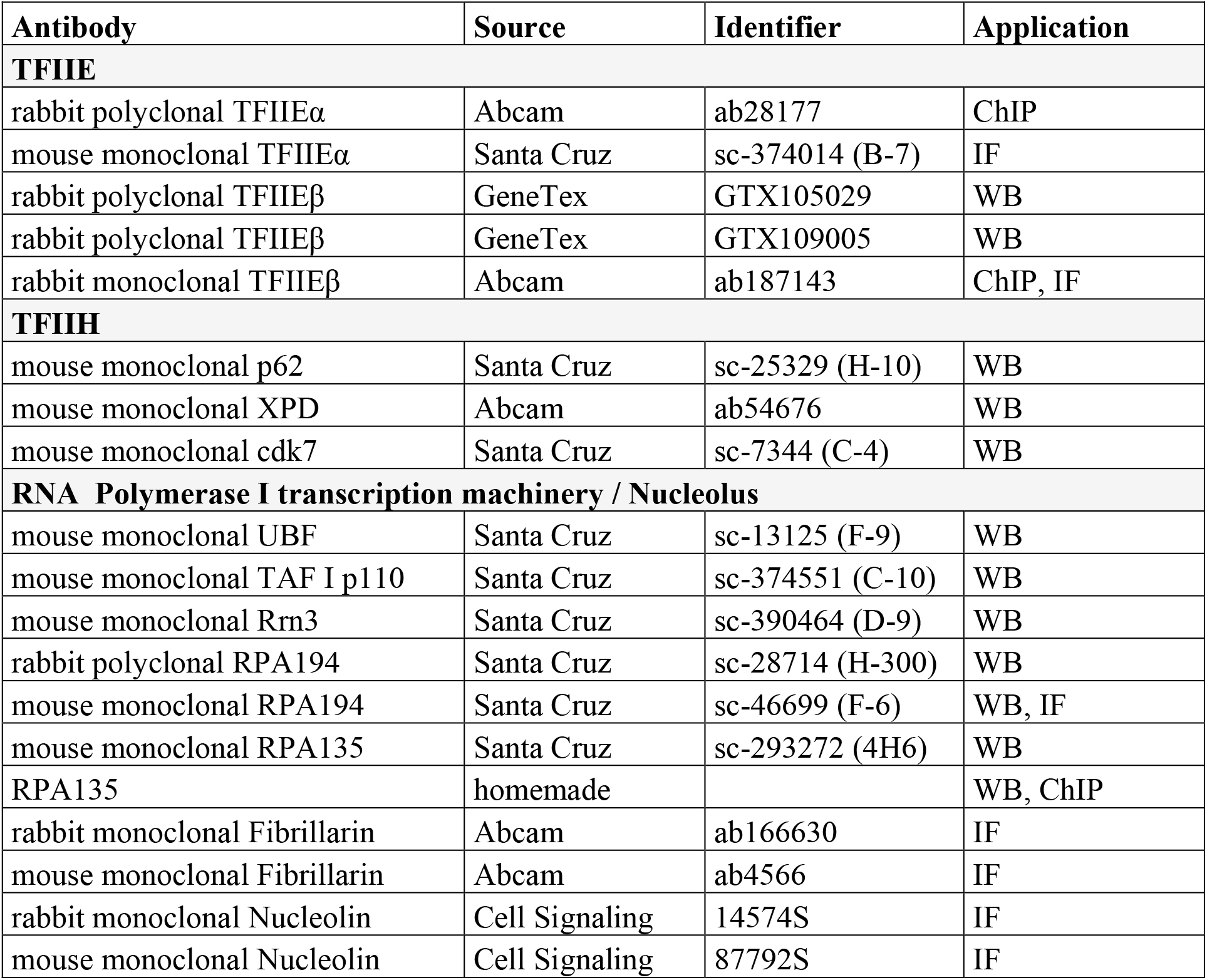
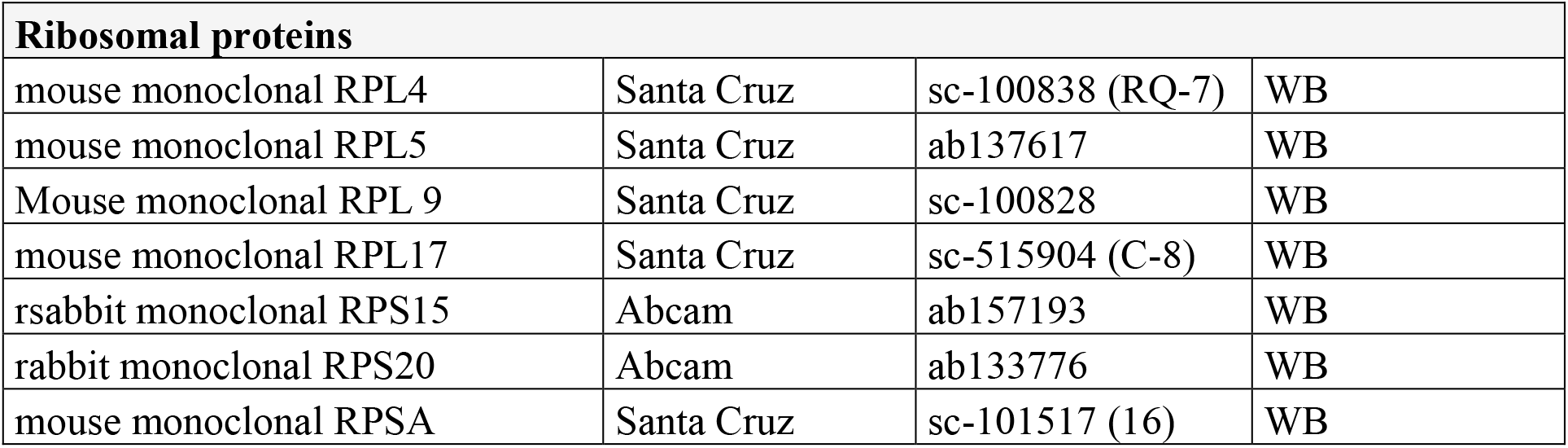
Antibodies of TFIIE, TFIIH, RNA polymerase I transcription machinery, Nucleolus and Ribosomal Proteins are listed below.

**Table S2.**
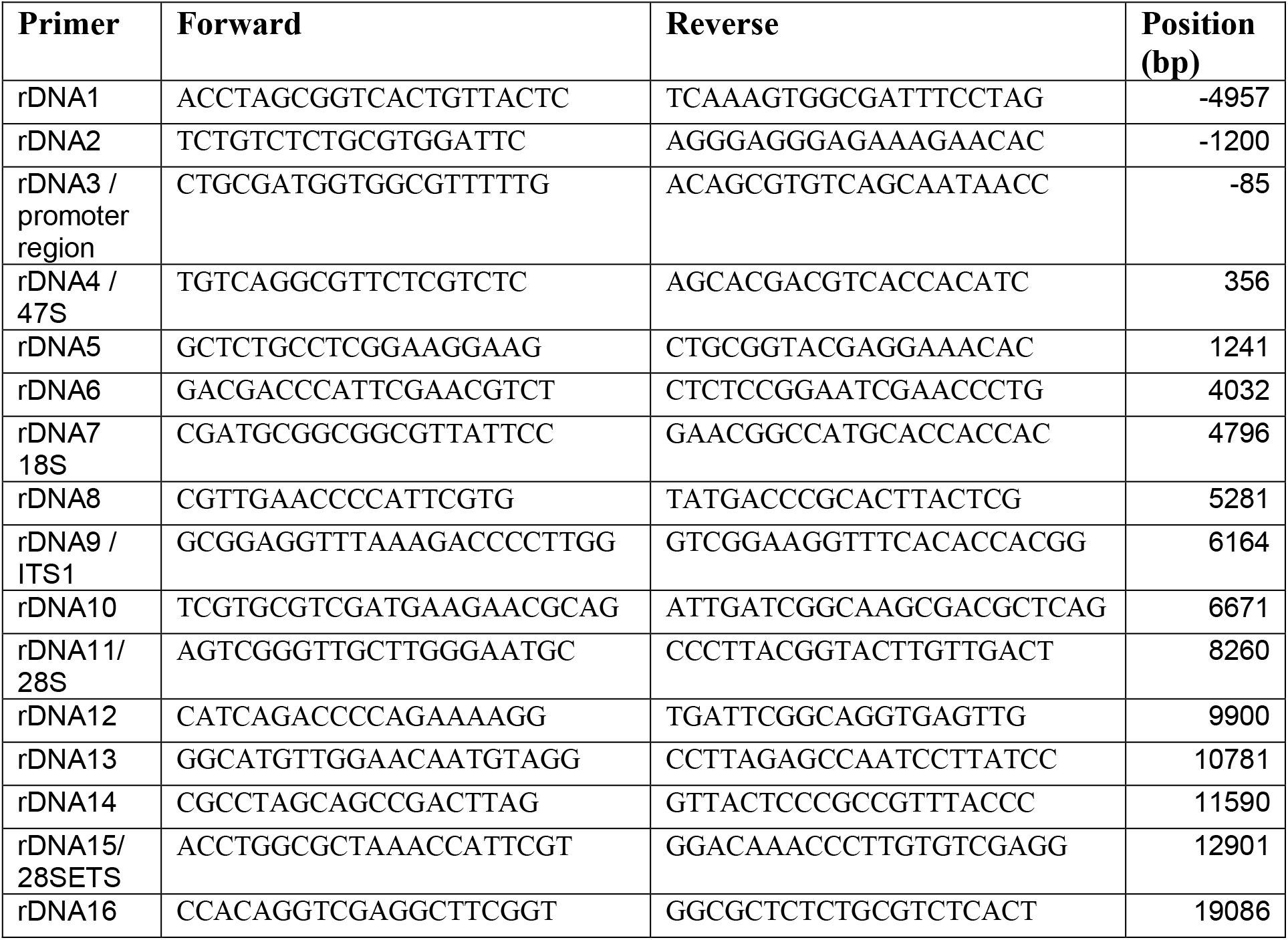
Sequences (5’ to 3’) of primers used to amplify rDNA regions are listed below.

